# *CAN OF SPINACH*, a novel long non-coding RNA, affects iron deficiency responses in *Arabidopsis thaliana*

**DOI:** 10.1101/2022.07.28.501865

**Authors:** Ahmet Bakirbas, Elsbeth L. Walker

**Affiliations:** Plant Biology Graduate Program, Department of Biology, University of Massachusetts Amherst, Amherst, MA, USA; Department of Biology, University of Massachusetts Amherst, Amherst, MA, USA

**Keywords:** long non-coding RNA, iron deficiency, nutrient deficiency, Arabidopsis thaliana, oxidative stress, singlet oxygen

## Abstract

Long non-coding RNAs (lncRNAs) are RNA molecules with functions independent of any protein-coding potential. A whole transcriptome (RNA-seq) study of Arabidopsis shoots under iron sufficient and deficient conditions was carried out to determine the genes that are iron-regulated in the shoots. We identified two previously unannotated transcripts on chromosome 1 that are significantly iron-regulated. We have called this iron-regulated lncRNA, *CAN OF SPINACH* (*COS*). *cos* mutants have altered iron levels in leaves and seeds. Despite the low iron levels in the leaves, *cos* mutants have higher chlorophyll levels than WT plants. Moreover, *cos* mutants have abnormal development during iron deficiency. Roots of *cos* mutants are longer than those of WT plants, when grown on iron deficient medium. In addition, *cos* mutant plants accumulate singlet oxygen during iron deficiency. The mechanism through which *COS* affects iron deficiency responses is unclear, but small regions of sequence similarity to several genes involved in iron deficiency responses occur in *COS*, and small RNAs from these regions have been detected. We hypothesize that *COS* is required for normal adaptation to iron deficiency conditions.

## 1 Introduction

Iron is an essential micronutrient for plant development and is required in many enzymatic processes such as the electron transport chain of respiration and photosynthesis, and heme biosynthesis (Tanaka et al., 2011; Connorton et al., 2017). All organisms strictly regulate iron uptake to ensure that only the required amount of iron is present in cells and tissues (Ravet et al., 2009). For these reasons, plants have developed complex mechanisms to tightly regulate iron uptake, use and storage while adapting to changes in iron concentration in the environment.

As a transition metal, iron is redox active (Ward and Crichton, 2016). Iron deficiency decreases photochemical capacity and photosynthetic pigments (*e*.*g*., chlorophyll) by diminishing the number of photosynthetic units per unit leaf area (Spiller and Terry, 1980). Because iron deficiency decreases the concentration of photosynthetic pigments, it can promote photoinhibition (Godde and Hefer, 1994). This impairment of photosynthetic electron transport can result in accumulation of reactive oxygen species (ROS) (Yadavalli et al., 2012; Tewari et al., 2013). In order to maintain photosynthesis, plants must respond to changes in iron status without producing high levels of ROS (Kroh and Pilon, 2020). In the reaction center of photosystem II, the main ROS produced is singlet oxygen (Krieger-Liszkay, 2005). Aberrant production of ROS leads to photooxidative damage and eventually phytotoxicity in the organism (Halliwell and Gutteridge, 2015). Among all the different ROS, singlet oxygen is the major species causing photooxidative damage in plants (Triantaphylidès et al., 2008).

Plants respond to Fe-deficiency by enhancing their iron uptake capacity and iron use efficiency, but also by altering development. Iron is essential for chlorophyll biosynthesis and during iron-deficiency, the chlorophyll content of leaves decreases, a process known as iron chlorosis (Msilini et al., 2013). Under Fe-deficiency, plants display low levels of iron in leaves and later in development, in seeds (Waters et al., 2006). Moreover, during Fe-deficiency, roots are shorter compared to plants grown under Fe-sufficient conditions (Li et al., 2016; Zanin et al., 2017). Plant yield and quality is reduced during iron deficiency and this presumed to be the result of decreased photosynthesis associated with iron chlorosis (Zhang et al., 2019).

In response to Fe-deficiency, many changes occur at the transcriptional level, and this has been particularly well studied in roots, where the network of genes involved in regulating iron uptake has been defined. Expression of *IRT1, FRO2* and a cascade of bHLH transcription factors are up-regulated in roots to activate the iron uptake machinery (Liang, 2022). While transcriptional response of the roots to Fe-deficiency is well studied, less is known about transcriptional changes occurring in leaves. Leaf responses to iron deficiency are rapid, and may occur even more quickly than responses in roots (Khan et al., 2018). Expression of *HEMA1* and *NYC1*, which are genes involved in photosynthesis and tetrapyrrole biosynthesis, respectively, were downregulated in Fe-deficient leaves (Rodríguez-Celma et al., 2013). The role of transcription factor POPEYE (PYE) is well characterized in the roots as a negative regulator of iron uptake and homeostasis (Long et al., 2010). In a recent study, *PYE* was shown to be upregulated in leaves under Fe-deficiency, and to lessen the potential for photooxidative damage in leaves (Akmakjian et al., 2021) The Fe-S cluster transfer protein NEET has critical roles in iron homeostasis in leaves (Nechushtai et al., 2012). Disruption of the single Arabidopsis NEET gene triggers Fe-deficiency responses that result in iron over-accumulation in chloroplasts and enhanced accumulation in leaves (Zandalinas et al., 2020).

Under limited nutrient conditions plants adjust their metabolism to tolerate limited nutrient quantities. Nutrient economy strategies are used by plants in these situations to prioritize, recycle and remobilize limiting nutrients (Blaby-Haas and Merchant, 2013). Iron economy strategies are initiated when shoot demand exceeds supply by the roots (Urzica et al., 2012; Blaby-Haas and Merchant, 2013; Hantzis et al., 2018). Under iron limited conditions, limited amounts of iron are preferentially allocated to the most important functions. Specific changes occur at the transcript and protein level as part of iron economy strategy to help adjust the metabolism (Hantzis et al., 2018).

Long non-coding RNAs (lncRNAs) are a poorly understood, but common feature in eukaryotic genomes. The definition of a lncRNA comprises three criteria: the RNA molecule must have a function, to distinguish it from random products of transcriptional noise; this function must be independent of any protein-coding capacity of the RNA; and a lncRNA must be generated through mechanisms different from those that produce sRNAs (*e*.*g*., Dicer endonucleases) (Wierzbicki et al., 2021). lncRNAs vary in size and structure and can affect gene expression of their targets by modifying chromatin accessibility, modifying transcription itself; affecting splicing processes or affecting translation. lncRNAs can act locally (*in cis*) to regulate genes near the locus that encodes them, or at a distance (*in trans*) to affect expression of genes that are located in other parts of the genome (Chen et al., 2021; Wierzbicki et al., 2021). lncRNAs appear to be capable of producing sRNA through the action of Dicer (Ma et al., 2014). Such sRNA could also potentially account for biological activities of lncRNA in trans.

In this study, we have identified a previously unannotated lncRNA gene that we named *CAN OF SPINACH* (*COS*). *COS* expression is regulated by Fe-deficiency in both shoots and roots. Two *cos* mutant alleles were identified, and both had low levels of iron in shoots and seeds. Surprisingly, despite low iron levels, *cos-2* shoots had increased levels of chlorophyll. Under Fe-deficiency conditions, *cos* mutants displayed abnormal growth. Also, cos mutants accumulate high levels of singlet oxygen in shoots, and appear to fail to make normal adaptations to low iron conditions. This work indicates that the previously unannotated lncRNA, *COS*, has a role iron homeostasis.

## 2 Materials and Methods

### Plant material and growth

The wild type *Arabidopsis thaliana* accession Columbia-0 (Col-0) was used for all experiments; T-DNA insertion lines for *COS* (SAIL_445_G04; *cos-1* and SALK_024945C; *cos-2*) were obtained from the Arabidopsis Biological Resource Center (ABRC). Homozygous mutant lines were confirmed by PCR. For soil growth, seeds were planted directly onto Pro-mix potting soil pre-treated with Gnatrol (Valent Bioscience Corporation) and stratified for three days at 4°C. Growth chamber conditions for soil-grown plants were 16 hours light (110 μmol·m^−2^ · s^−1^), 8 hours dark at 22°C. For growth in sterile culture, sterilized seeds were plated directly on sterile petri plates containing 1/2X Murashige and Skoog medium (1/2X MS+Fe, prepared with 10X macronutrient solution, 100X micronutrient solution (without iron) and 100X Ferrous sulfate/chelate solution from PhytoTech Labs, Inc.) and placed at 4°C for three days. 1/2X MS-Fe was prepared using 10X macronutrient solution and 100X micronutrient solution (without iron). Growth chamber conditions for plate-grown plants were 16 hours light (110 μmol·m^−2^ · s^−1^), 8 hours dark at 22°C.

### RNA extraction

Root and shoot tissues were disrupted in 1.5 mL Eppendorf tubes using a tissue homogenizer (Qiagen TissueLyser II) with 3.2 mm chrome steel beads (BioSpec Products). Total RNA was isolated using the Direct-zol(tm) RNA MiniPrep Kit (Zymo Research) with DNase I treatment according to the manufacturer’s instructions.

### cDNA synthesis, RT-PCR and qRT-PCR

cDNA was synthesized from total RNA using SuperScript IV VILO Master Mix (Thermo Fisher Scientific) according to the manufacturer’s instructions. The RT-PCR and qRT-PCR reactions were carried out as described previously (Kumar et al., 2017). The primers used in this study are given in Supplementary Table 1.

### RNA-seq and data analysis

Three biological replicates of three conditions were used for RNA-seq analysis. +Fe samples: plants were grown on 1/2X MS+ Fe for 14 days, then were transferred to fresh 1/2X MS+Fe for 3 days, and finally were transferred to 1/2X MS+Fe for 24 hours; -Fe samples: plants were grown on 1/2X MS+ Fe for 14 days, then were transferred to 1/2X MS-Fe for 3 days, and finally were transferred to fresh 1/2X MS-Fe for 24 hours; resupply samples: plants were grown on 1/2X MS+ Fe for 14 days, then were transferred to 1/2X MS-Fe for 3 days, and finally were transferred to 1/2X MS+Fe for 24 hours. Paired-end libraries were generated and sequenced by Novogene (Sacramento, CA).

Adapter sequences and low-quality reads were removed from raw reads with Trimmomatic v0.36.5 (Bolger et al., 2014). Mapping of reads to the Arabidopsis TAIR10 genome was carried out using HISAT2 v2.1.0 (Kim et al., 2015). Reads were assembled into transcripts and quantified using StringTie v1.3.3.2 (Pertea et al., 2015). Annotation was done using the TAIR10 genome GTF annotation file (www.arabidopsis.org). Differential gene expression analysis was performed using edgeR v3.20.7.2 through the Galaxy platform (Afgan et al., 2018). Minimum log2 fold change of 1 and Benjamini-Hochberg adjusted p-value of 0.05 were employed (Benjamini and Hochberg, 1995) as cutoff values.

### Root length measurements

Col-0, *cos-1* and *cos-2* seeds were grown on 1/2X MS+Fe plates for seven days and shifted to either 1/2X MS+Fe or 1/2X MS-Fe plates for three days. Plates were scanned using a flatbed scanner (Epson V700), and root length was measured using ImageJ software v1.52 (Schindelin et al., 2012).

### Chlorophyll determination

Plants were grown directly on soil. Shoot tissues were collected when the first inflorescence became visible within the rosette, and fresh weight was determined. Chlorophyll extraction and quantification were performed as described previously (Waters et al., 2006).

### Iron content determination by ICP-MS

Arabidopsis seeds for Col-0, *cos-1* and *cos-2* were grown directly on soil to determine iron content in shoots. Whole rosettes were collected when the first inflorescence became visible within the rosette and dried at 50°C for three days. To determine iron content in seeds, plants for Col-0, *cos-1* and *cos-2* were grown directly on soil until the seeds were mature. Seeds from individual plants were collected in Eppendorf tubes. ICP-MS analysis was carried out at the Ionomics Facility at the Donald Danforth Plant Science Center.

### Seed weight determination

Seed weight was determined based on the weight of 10 batches of 100 seeds from three individual plants for each genotype. Weight per seed was calculated from the average of 10 seed batches. Seed number was determined by dividing the total weight of seeds by the individual seed weight.

### Singlet oxygen detection

Plants were grown on 1/2X MS+Fe plates for 14 days and shifted to either 1/2X MS+Fe or 1/2X MS-Fe plates for three days. The singlet oxygen sensor green (SOSG) staining procedure was performed as described by Akmakjian et al. (2021). Imaging of singlet oxygen detection was done on a Nikon MZ16 FA stereomicroscope equipped with a mercury lamp and a color camera.

### sRNA-seq and data analysis

Three biological replicates of three conditions were used for sRNA-seq analysis. +Fe samples: plants were grown on 1/2X MS+ Fe for 14 days, then were transferred to fresh 1/2X MS+Fe for 3 days, and finally were transferred to 1/2X MS+Fe for 24 hours; -Fe samples: plants were grown on 1/2X MS+ Fe for 14 days, then were transferred to 1/2X MS-Fe for 3 days, and finally were transferred to fresh 1/2X MS-Fe for 24 hours; resupply samples: plants were grown on 1/2X MS+ Fe for 14 days, then were transferred to 1/2X MS-Fe for 3 days, and finally were transferred to 1/2X MS+Fe for 24 hours.

Libraries were generated and sequenced by Novogene (Sacramento, CA). Cutadapt was used to remove adapter sequences (Martin, 2011). Shortstack was used to align, annotate, and quantify small RNAs (Axtell, 2013; Shahid and Axtell, 2014). The search size parameter for discovering sRNA loci with Shortstack was between 17 and 29 nt long. After processing the raw reads, raw read counts were used as input for differential expression analysis with DESeq2 (FDR < 10% and Log2FC ≥ |0.5|) (Love et al., 2014). Arabidopsis sRNA Target Prediction webserver at Donald Danforth Plant Science Center (wasabi.ddpsc.org/∼apps/tp/) was used to predict the targets of the iron up-regulated sRNA candidates. The default scoring system was used for the analysis.

## 3 Results

### Discovery of a Novel Iron-Regulated Long Noncoding RNA in Arabidopsis

A whole transcriptome (RNAseq) study of Arabidopsis shoots under iron sufficient and deficient conditions was carried out to determine which genes are iron-regulated in the shoots. Among the differentially expressed genes, a particular locus containing three protein-coding genes (AT1G13607, AT1G13608, AT1G13609) was scored as having iron-deficiency-regulated gene expression. Two of the genes (AT1G13608, AT1G13609) appeared to be upregulated during iron deficiency, and these genes have been identified as iron-regulated in other analyses (Rodríguez-Celma et al., 2013; Naranjo-Arcos et al., 2017; Bastow et al., 2018; Nam et al., 2021; Peixoto et al., 2021). Upon closer inspection of the count data (FPKM values), we noticed that the iron-regulated reads were mostly not coming from the strand encoding AT1G13607, AT1G13608, and AT1G13609. Instead, we identified a previously unannotated transcript on the reverse strand that is significantly iron-regulated (Fig. 1A and B). The AT1G13609 transcript on the top strand appears to be iron-regulated (Fig. 1B) but the difference in FPKM was not significant. Moreover, the FPKM values (89.7 and 55.3) indicate that the unannotated bottom strand transcript is expressed at much higher levels than the top strand genes AT1G13607, AT1G13608, and AT1G13609 (0.00001, 0.000012, 1.68 and 34.1)

**Figure 1.**
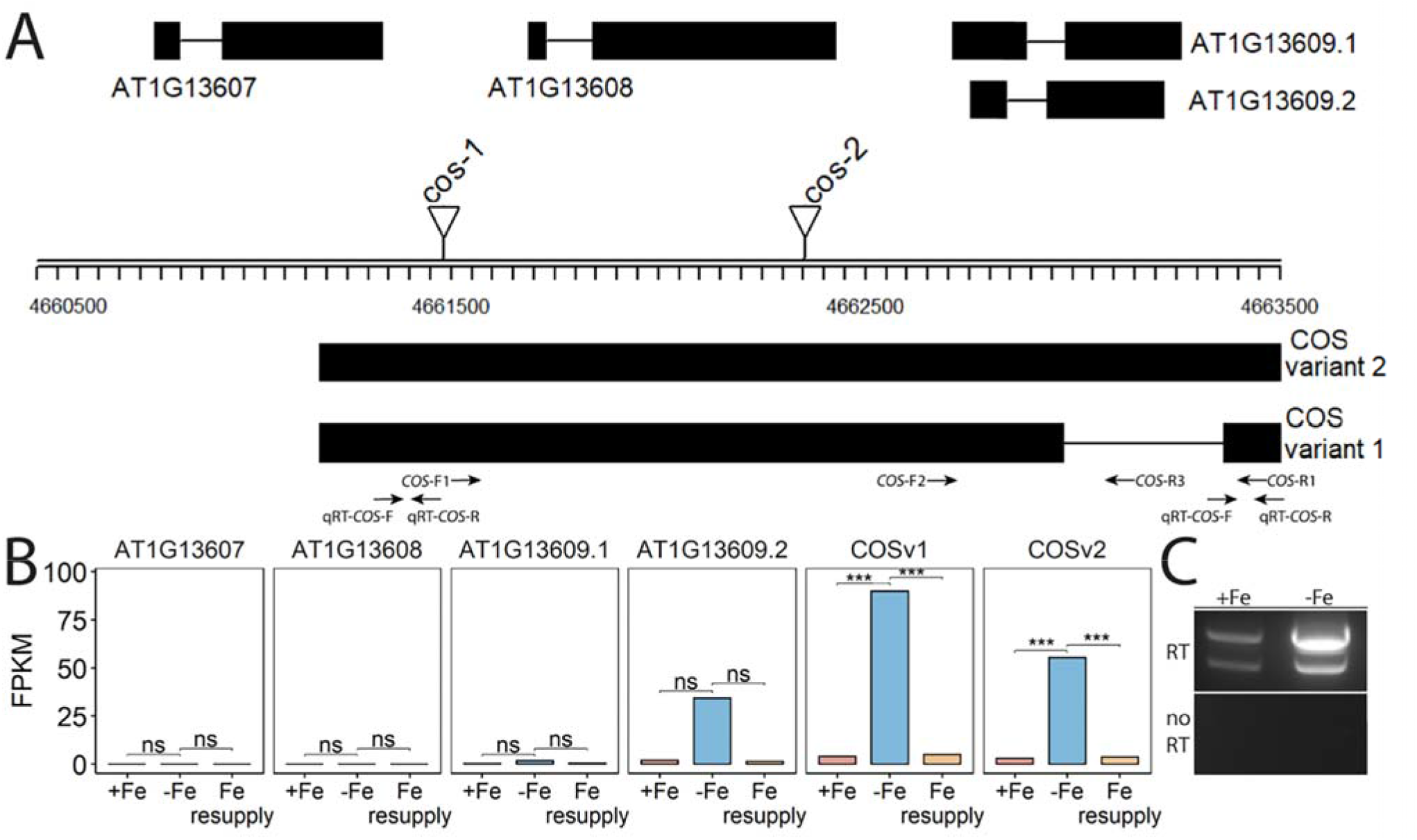
*CAN OF SPINACH* (*COS*), a novel long non-coding RNA (lncRNA). (A) Schematic representation of the locus. On the top strand, three genes (AT1G13607, AT1G13608 and AT1G13609) are annotated. The lncRNA *COS* is on the bottom strand. Thick boxes represent exons, lines represent introns. Positions of T-DNA insertion mutants used in this study are represented as triangles. Primers used in the study are represented as arrows at the bottom. (B) Bar plot representing the expression levels of various predicted transcripts under Fe sufficient, Fe deficient and Fe resupply conditions. Values from RNAseq are represented as fragments per kilobase of exon per million mapped fragments (FPKM). (C) RT-PCR using primers COS-F1 and COS-R1, using total RNA from shoots. Absence of genomic DNA contamination was confirmed by the lack of amplification of th no-RT control. cDNA was synthesized using oligodT_(20)_. *** represents p-value ≤ 0.0001. ns. not significant.

Using primers located at the 5’ and 3’ ends of the putative transcript (COS-F1 and COS-R1), we were able to demonstrate the existence of the full length, spliced transcript using reverse transcription PCR (RT-PCR). These primers were specifically designed to only detect the expression of the putative transcript and not the coding genes on the top strand (Fig. 1A). Both a spliced and an unspliced variant were amplified. Because cDNA synthesis was primed with oligodT, the transcript appears to be polyadenylated, as most plant lncRNAs are (Wang and Chekanova, 2017) (Fig. 1C). We named the newly identified transcript, *CAN OF SPINACH* (*COS*). We carried out an *ab initio* prediction of any peptides *COS* might encode. We found that *COS* has the potential to encode only a small peptide of 57 aa long. Considering the small size of this peptide relative to the length of the transcript, *COS* appears to be a long non-coding RNA (lncRNA; Wierzbicki et al., 2021).

Two different mutant alleles, *cos-1* (SAIL_445_G04) and *cos-2* (SALK_024945), in which T-DNAs are inserted in a *COS* exon (Fig. 1A), were obtained, and homozygous mutant plants were established. We performed RT-PCR using two sets of primers, one upstream of the insertion sites and one downstream of the insertion sites. By sampling different parts of the COS lncRNA, we observed that the cos transcripts in the mutants are likely to be aberrant. Specifically, RT-PCR analysis of the *COS* transcript levels (Fig. 2A) indicates that *cos-1* has detectable RNA expression with both primer sets, but is detected at significantly lower levels with one of these sets. Thus, *cos-1* may be a partial loss of function allele. The *cos-2* allele has no detectable transcripts with one of the primer sets used and is therefore likely to be a total loss of function allele. It may produce an aberrant transcript that lacks a portion of the normal RNA. It is possible that *cos-1* affects AT1G13608, and that *cos-2* affects AT1G13609, on the top strand. The level of expression of these genes is undetectably low in RT-PCR in every condition we tested, and the genes appear not to be regulated by iron deficiency. Thus, although it is possible that the mutations also affect these genes the expression of *COS* is very obviously affected. Although *COS* was first identified in Arabidopsis shoots, it is also expressed in the roots. The levels of *COS* transcripts in shoots and roots were similar when plants were grown with sufficient iron. During Fe-deficiency, the amount of *COS* transcript in roots trends lower, but this was not significant at our stringent P ≤ 0.01 criterion (Fig. 2B).

**Figure 2.**
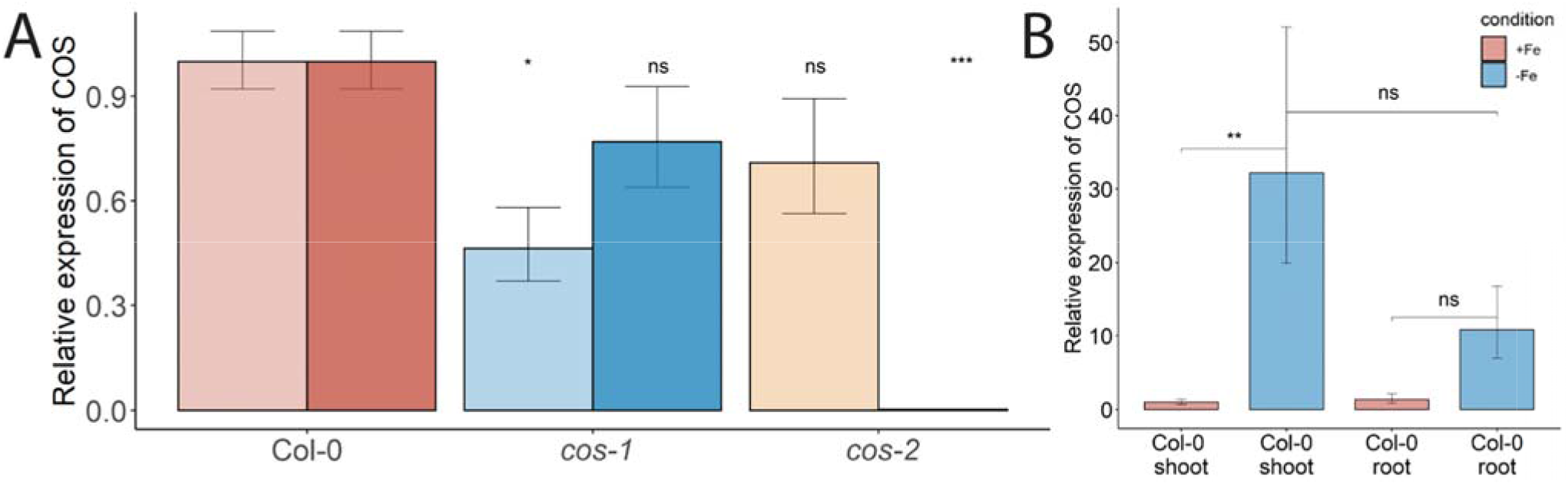
*COS* expression in WT and T-DNA mutants. (A) Two sets of primers were used for RT-PCR. The primer set upstream of the T-DNA insertion is designated with the lighter color, while the primer set downstream of the insertion is designated with darker color. Seedlings were grown on 1/2X MS agar with 50 μM Fe-EDTA for 5 d followed by a growth period of 3 d either on Fe-sufficient or Fe-deficient MS agar medium. Bars represent means ± SD of relative transcript levels normalized to *ACT2* (n=3). (B) Expression levels of *COS* transcript in WT shoots and roots. Seedlings were grown on 1/2X MS agar with 50 μM Fe-EDTA for 14 d followed by a growth period of 3 d either on Fe-sufficient or Fe-deficient MS agar medium. Bars represent means ± SD of relat ve transcript levels normalized to *ACT2* (n=3). * represents p-value ≤ 0.01. ** represents p-value ≤ 0.001, *** represents p-value ≤ 0.0001, ns. not significant.

### cos Mutants Have Decreased Fe Levels in Shoots and Seeds

Phenotypic characterization of T-DNA insertion mutants was carried out to further investigate the role of *COS* in iron homeostasis. We measured the accumulation of iron in the shoots of wild type (WT; Col-0) and *cos* mutant plants using ICP-MS (Fig. 3A). The *cos* mutants have significantly less iron in their shoots than WT plants. The levels of Zn, Cu, and Mn in the mutants were not significantly different from WT (not shown). To complement and confirm these ICP-MS findings, we examined the expression of *Ferritin 1* (*FER1*) in shoots. *FER1* mRNA levels are commonly used as a marker for shoot iron status (Gaymard et al., 1996). In accord with shoot iron levels, *FER1* expression was significantly lower in *cos-2* mutants, while *FER1* levels trended low, but were not significantly different in the partial loss of function allele *cos-1* (Fig. 3B). We postulated that the low iron levels in *cos* mutant plants might result in low iron levels in seeds. To examine this we analyzed iron accumulation in seeds. We found that seeds of *cos* mutants contain significantly less iron than WT seeds (Fig. 3C). The levels of other metals in cos mutants were not significantly different from WT (not shown).

**Figure 3.**
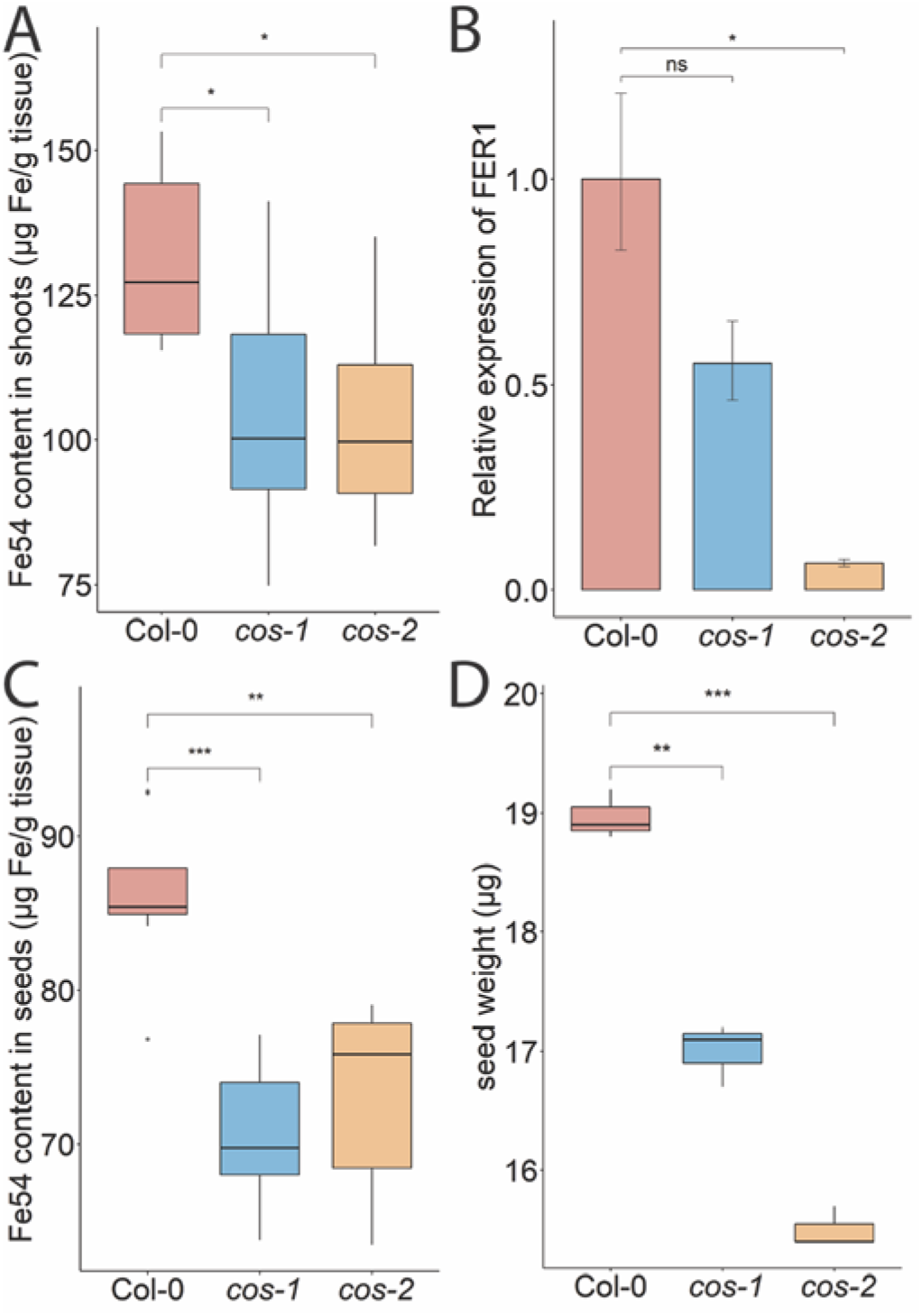
Phenotype of *cos* T-DNA mutants on soil. Iron concentration of shoots (A) and seeds (C) from plants grown on commercial potting mix. Results are given as μg Fe per gram of tissue (n=8). (B) Expression levels of *FER1* transcript in WT (Col-0) and *cos* mutants. Seedlings were grown on 1/2X MS agar with 50 μM Fe-EDTA for 14 d followed by a growth period of 3 d either on Fe-sufficient or Fe-deficient MS agar medium. Bars represent means ± SD of relative transcript levels normalized to *ACT2* (n=3). (D) Individual seed weight of WT and *cos* mutants was determined by sampling weights of 10 batches of 100 seeds. * represents p-value ≤ 0.01. ** represents p-value ≤ 0.001, *** represents p-value ≤ 0.0001, ns. not significant.

Since the iron levels in the *cos* seeds were low, we wondered whether this would affect seed development. We measured the seed weight and seed number in WT and *cos* mutants. The individual seed weight was decreased in *cos* mutants compared to WT seeds grown at the same time (Fig. 3D). Both *cos-1* and *cos-2* displayed upward trends in seed number compared to WT, but these differences were not statistically significant (not shown).

### Growth phenotypes of cos *Mutants*

One of the classic symptoms of iron deficiency is chlorosis (low chlorophyll), and many mutants that accumulate lower levels of iron in their tissues also exhibit iron deficiency chlorosis (Robinson et al., 1999; Curie et al., 2001; Vert et al., 2002; Colangelo and Guerinot, 2004; Waters et al., 2006). Because *cos* mutant shoots have low iron levels, we expected that they might have low chlorophyll levels. Surprisingly though, shoots of *cos-2* knockout plants had significantly more chlorophyll than WT. The chlorophyll content of *cos-1* mutants appeared to trend higher than WT but was not significantly higher (Fig. 4A). Under iron deficient conditions, *cos* mutants displayed chlorosis similar to WT, however, due to the low levels of chlorophyll during iron deficiency we were not able measure the -Fe chlorophyll levels reliably.

**Figure 4.**
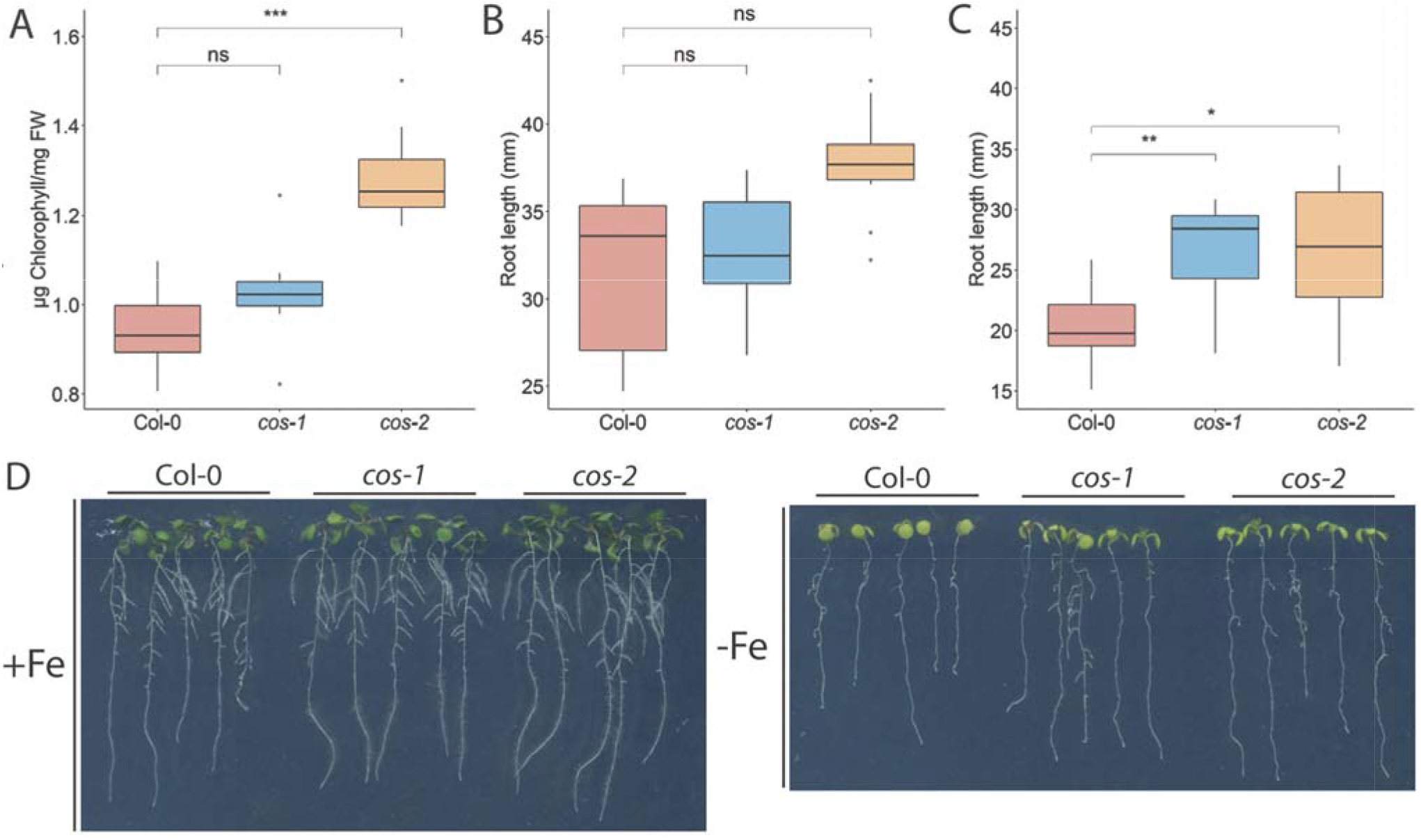
Phenotype of *cos* T-DNA mutants on 1/2X MS medium. (A) Total chlorophyll concentration of the shoot systems of plants grown on soil for 14 d. (B,C) Total root length of seedlings grown on 1/2X MS agar medium either with 50 μM Fe-EDTA (+Fe) or without Fe (-Fe) for 10 days. (D) Seedlings were grown on 1/2X MS agar medium either with 50 μM Fe-EDTA (+Fe) or without Fe (-Fe) for 10 days. * represents p-value ≤ 0.01, ** represents p-value ≤ 0.001, *** represents p-value ≤ 0.0001, ns. not significant.

We examined the root length of WT and *cos* mutants because we hypothesized that with low iron levels in shoots, *cos* mutants might show a root phenotype under Fe-deficiency. The root lengths of both *cos* mutants are similar to WT under iron sufficient conditions (Fig. 4B). However, upon shifting plants to Fe-deficient environment for three days, both *cos-1* and *cos-2* had significantly longer root lengths than WT plants (Fig. 4C).

Another symptom of iron deficiency is the up-regulation of transcripts for genes encoding the iron transporter, *IRT1*, and the iron-uptake regulatory gene *FIT* (Connolly et al., 2002; Colangelo and Guerinot, 2004). We examined *IRT1* and *FIT* expression in WT and *cos* roots under Fe-sufficient and deficient conditions. Expression of *IRT1* and *FIT* were similar to WT in *cos* roots under both conditions tested (Suppl. Fig. 1), in spite of the low iron levels measured in *cos* mutant shoots and seeds.

### cos *Mutants Accumulate Singlet Oxygen in Shoots*

Iron deficiency causes well documented negative effects on photosynthesis, and plants adapt to iron deficiency by decreasing the expression of particular iron containing proteins in leaves, while leaving the expression of presumably more essential iron proteins intact (Hantzis et al., 2018). During iron deficiency, plants produce reactive oxygen species, including singlet oxygen, and have specific responses that mitigate the damage that iron deficiency can cause in leaves (Akmakjian et al., 2021). Specifically, *bHLH105*/*ILR3* and *PYE* were shown to be required for photoprotection during Fe-deficiency and prevent accumulation of singlet oxygen (Akmakjian et al., 2021). We wondered whether disruption of COS affected these responses.

We investigated whether ROS production was affected in *cos* mutants. Expression of *OXI1*, an oxidative stress marker whose expression is induced by both singlet oxygen and hydrogen peroxide (Rentel et al., 2004), was significantly increased in *cos* mutants compared to WT under Fe-deficiency (Fig. 5A). We then checked the expression of *AT3G03810*, a gene whose expression is induced specifically by singlet oxygen (Koh et al., 2016). During Fe-deficiency, *AT3G03810* expression was significantly increased in *cos* mutants compared to WT (Fig. 5B). The rise in expression of both of these genes during iron deficiency suggests that *cos* mutants might be accumulating singlet oxygen during Fe-deficiency.

**Figure 5.**
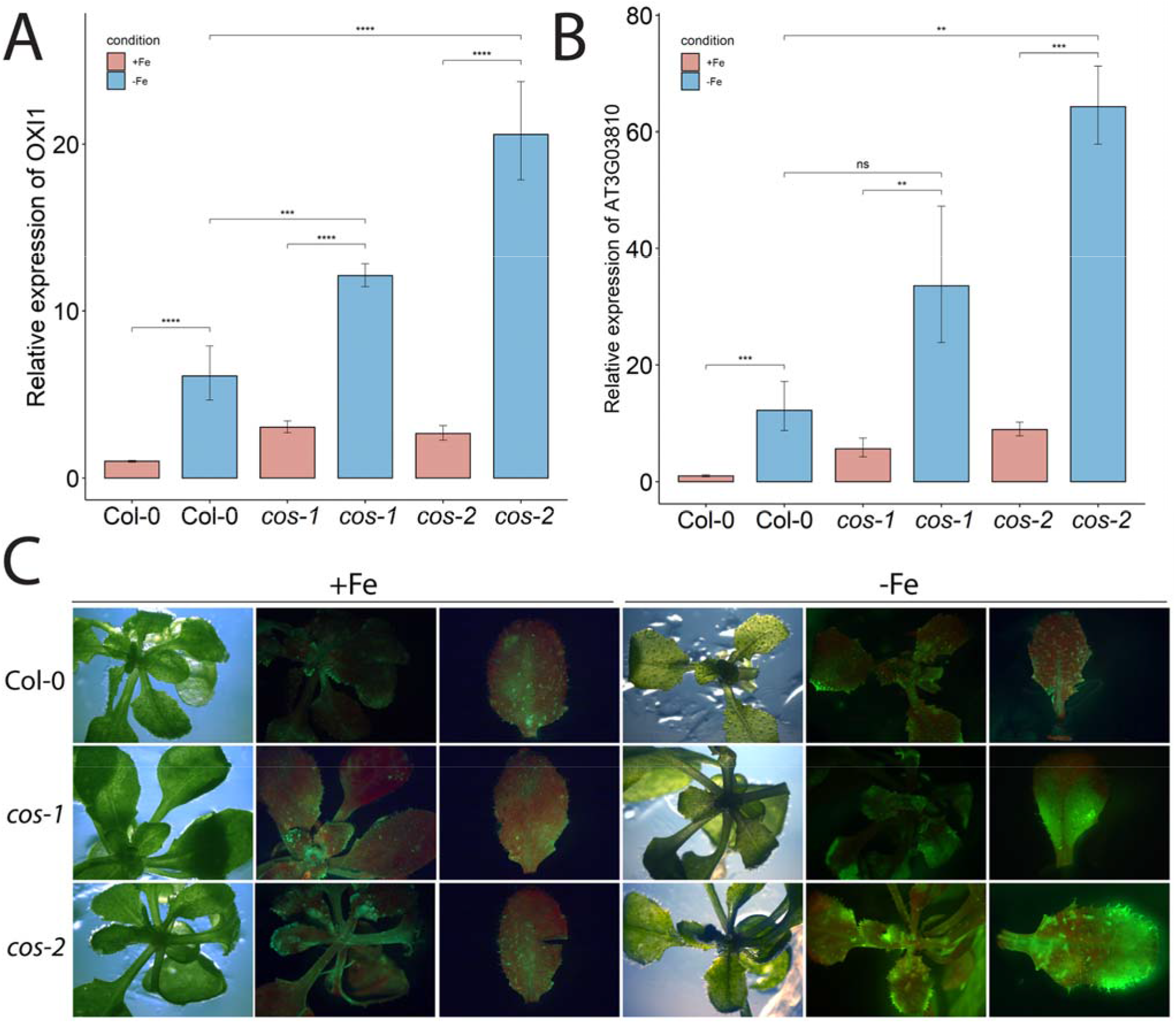
ROS in leaves of *cos* T-DNA mutants. RT-PCR of oxidative stress marker *OXI*1 (A) and singlet oxygen marker *AT3G0381*0 (B) in WT and *cos* mutant shoots. Seedlings were grown on 1/2X MS agar with 50 μM Fe-EDTA for 14 d followed by a growth period of 3 d either on Fe-sufficient or Fe-deficient MS agar medium. Bars represent means ± SD of relative transcript levels normalized to ACT2 (n=3). (C) Singlet oxygen detection in *cos* mutants by SOSG. Seedlings were grown on 1/2X MS agar with 50 μM Fe-EDTA for 14 d followed by a growth period of 3 d either on Fe-sufficient or Fe-deficient MS agar medium. Seedlings were stained with SOSG and incubated under high light (500 µmol photons m^-2^ s^-1^) for 30 minutes. SOSG fluorescence is represented by green while chlorophyll autofluorescence is represented by red. * represents p-value ≤ 0.01, ** represents p-value ≤ 0.001, *** represents p-value ≤ 0.0001, ns. not significant.

Next, we used the fluorescent stain Singlet Oxygen Sensor Green (SOSG; Invitrogen) to detect singlet oxygen production in *cos* mutants. Seedlings were vacuum infiltrated in the dark with SOSG, followed by a brief incubation under high light and subsequently imaged using a stereomicroscope. In WT plants, fluorescence from SOSG was barely detectable in either Fe-sufficient or deficient conditions. No fluorescence was detected in control plants that were infiltrated with buffer alone (Suppl. Fig. 2). For the *cos* mutants, the fluorescence was similar to WT in Fe-sufficient conditions, but markedly brighter when plants were grown without iron, indicating that *cos* mutants accumulate high levels of singlet oxygen during Fe-deficiency (Fig. 5C).

### COS *has small regions of sequence complementarity with several iron deficiency genes*

While analyzing the *COS* locus, we noticed an approximately 600 bp region within *COS* that had small (17-20 nt) sites of sequence complementarity to several well-known Fe-deficiency response genes (*bHLH38, bHLH39, bHLH100, bHLH101, bHLH105/ILR3, bHLH115, BTSL1, FSD1, IMA1, PYE* and *VIT1*; Fig. 6; Liang, 2022). *COS* also contains two overlapping sites with sequence complementarity for the *PHT4;4* gene, which is required for phosphate-deficiency induced inhibition of iron chlorosis (Nam et al., 2021; Fig. 6). Since lncRNAs can produce sRNA, apparently through Dicer activity (Ma et al., 2014), we performed small RNA sequencing (sRNAseq) data from iron deficient and iron sufficient shoots, and discovered that there are small RNAs (sRNAs) generated, apparently from the larger *COS* transcript, that might target these Fe-deficiency genes, ultimately affecting their expression. The number of sRNAs emanating from COS is markedly increased during iron deficiency (Fig. 6)

**Figure 6.**
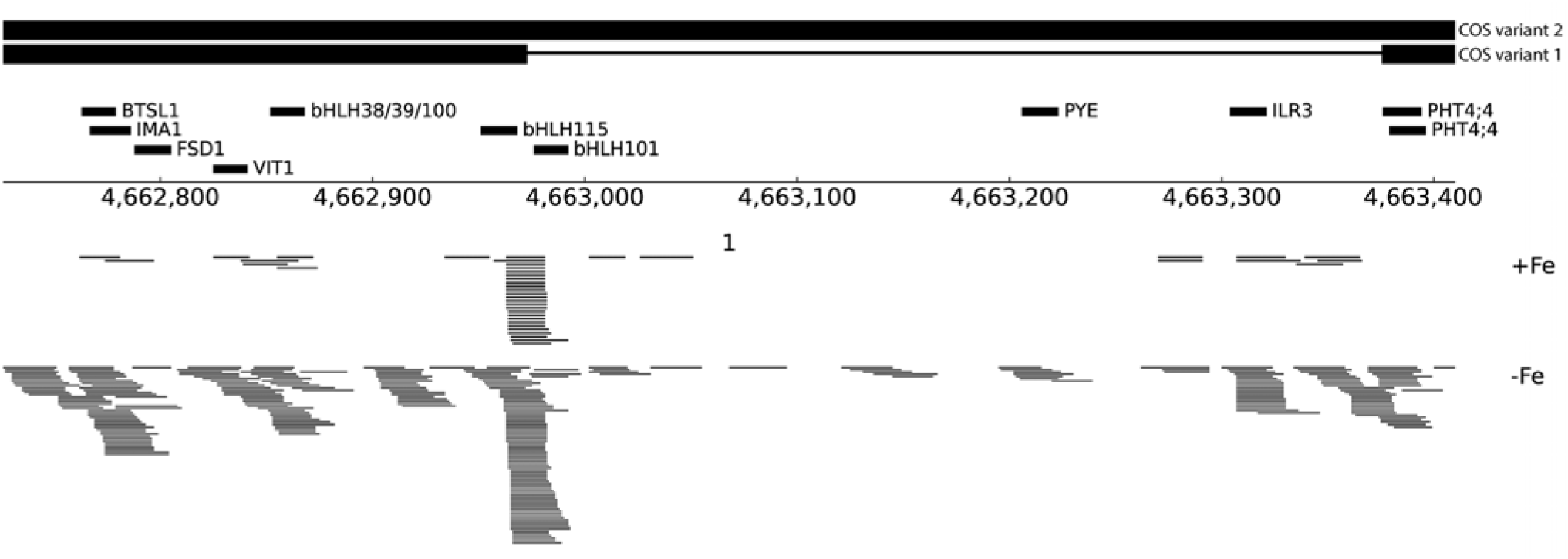
Small RNA reads at the *COS* locus. (A) Schematic representation of an ∼600 bp region within *COS* that shares sequence complementarity to several Fe-deficiency genes, indicated with thick black lines labeled with putative target genes. At bottom, sRNAseq reads mapping to COS are shown from both +Fe and -Fe samples.

We checked the expression of some of these targets to test if their expression levels are affected in *cos* mutants. *bHLH105*/*ILR3* expression, like the expression of other bHLH subgroup IVc transcription factors, is not affected by Fe-deficiency (Li et al., 2016), but expression of the *PYE* gene increases during iron deficiency (Rodríguez-Celma et al., 2013). We observed that *cos* mutant shoots failed to induce *PYE* expression under Fe-deficiency, while *PYE* expression in WT increased almost three-fold in iron-sufficient leaves (Fig. 7A). This could imply that normal PYE functions are affected in *cos* mutant plants. We did not detect any changes in *ILR3* expression under Fe-deficient conditions compared to its expression in WT. Under Fe-sufficient conditions, *ILR3* expression was significantly decreased in *cos* mutants compared to WT (Fig. 7C). While expression levels of *IMA1* and *BTSL1* were not different than it is in WT under Fe-sufficient conditions, the difference in their expression compared to WT was more pronounced under Fe-deficiency (Fig. 7D and E). We did not see any difference in expression of *bHLH38* in *cos* mutant shoots under Fe-sufficient or deficient conditions (Fig. 7F).

**Figure 7.**
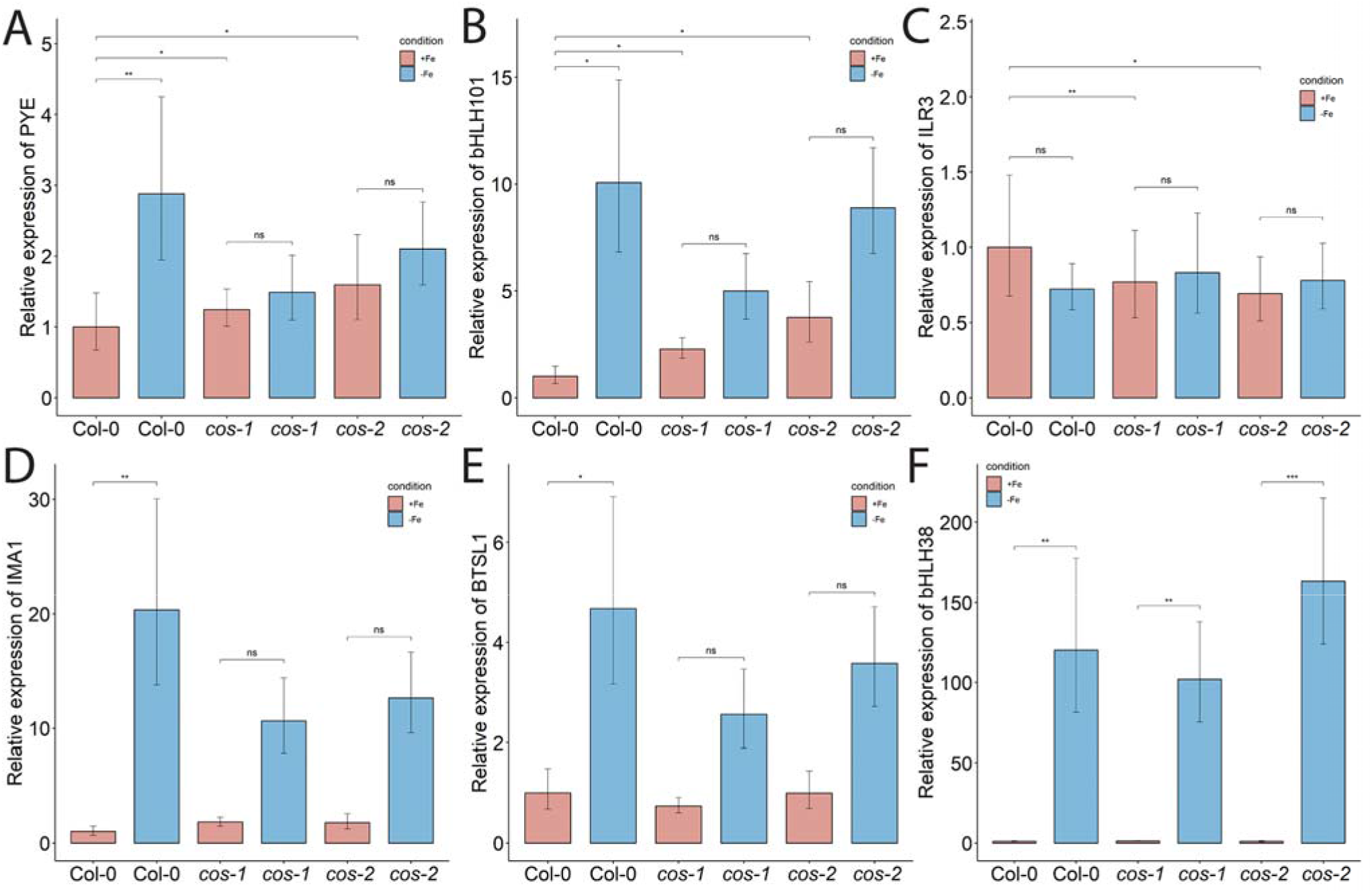
Expression of iron deficiency regulated genes in *cos* mutants. Expression levels of *PYE* (A), *bHLH101* (B), *ILR3/bHLH105* (C), *IMA1* (D), *BTSL1* (E), and *bHLH38* (F) transcripts in WT shoots and roots. Seedlings were grown on 1/2X MS agar with 50 μM Fe-EDTA for 14 d followed by a growth period of 3 d either on Fe-sufficient or Fe-deficient MS agar medium. Bars represent means ± SD of relative transcript levels normalized to *ACT2* (n=3). * represents p-value ≤ 0.01. ** represents p-value ≤ 0.001, *** represents p-value ≤ 0.0001, ns. not significant.

## 4 Discussion

In this study, we have identified a previously unannotated iron-regulated long non-coding RNA gene that we named *CAN OF SPINACH* (*COS*). It is possible that previous gene expression studies could have misidentified the genes on the opposite strand (AT1G13608 and/or AT1G13609) as iron-regulated, and that the truly iron-regulated mRNA from the locus is actually the *COS* lncRNA (Rodríguez-Celma et al., 2013; Naranjo-Arcos et al., 2017; Bastow et al., 2018; Nam et al., 2021; Peixoto et al., 2021). We detected a spliced and an unspliced variant of *COS* in whole shoots of Arabidopsis, both of which were polyadenylated (Fig. 1C).

*cos* mutant plants have phenotypes that suggest abnormalities in iron homeostasis. They have reduced levels of iron in their shoots and seeds, which indicates that they are not taking up normal amounts of iron from the soil. In addition, and contrary to the expectation for a plant with lowered iron in tissues, the mutants have elevated chlorophyll levels (Fig. 4A). This indicates that the plants are not responding to low shoot iron levels in a typical way. The roots of *cos* mutants also have abnormal growth, albeit only during iron deficiency. The *cos* roots are long compared to WT roots when they are grown without iron in the medium (Fig. 4B and C). This phenotype can be interpreted in two ways. In one view, when iron is not available, root growth simply cannot continue owing to the lack of iron *per se*. However, it is possible that reduced root growth during iron deficiency is a low-iron-induced developmental program intended to protect the plant and reserve accumulated iron for more critical functions. The high chlorophyll/low iron phenotype of the mutant could be interpreted in a similar way: WT plants may reduce chlorophyll levels as an iron economy measure, when iron in the shoots is low. In *cos* mutants, that iron economy measure may have been disrupted, and the mutants produce high amounts of chlorophyll in spite of the low levels of iron in the shoots. If this view is correct, then the *cos* phenotypes could be interpreted as a failure of the plant to make normal adaptive growth alterations during iron deficiency. Since *cos* mutants seem to fail to adapt normally to iron, we considered whether *cos* mutants might have a reproductive penalty, and they do. The individual seed weight is lower in both *cos* mutants (Fig. 3D), which could indicate minor problems with seed development. However, seed number was not strongly affected in the mutants.

Our singlet oxygen findings also support the idea that *cos* mutants do not adapt well to iron deficiency. One of the ways that Arabidopsis adapts to low iron is by changing gene expression to lessen the potential for photooxidative damage that occurs during iron deficiency (Akmakjian et al., 2021). The *cos* mutant plants are less able to prevent accumulation of singlet oxygen during iron deficiency, possibly owing in part to their inability to induce expression of *PYE*, and/or their lowered basal level of *BHLH105/ILR3*, which are both required for this adaptation (Fig. 7B). Increased gene expression from the generic oxidative stress marker *OXI1* and *At3g01830*, which is specifically induced by singlet oxygen, also indicate that *cos* plants have increased singlet oxygen (Fig. 5A and B.

The idea that *cos* mutants could fail to adapt to limited nutrient conditions brought the concept of nutrient economy strategies into mind. Nutrient economy strategies are present for several metals in a variety of organisms, and serve to prioritize, recycle and remobilize limiting nutrients (Blaby-Haas and Merchant, 2013). In plants, micronutrient economy strategies have been documented for iron, copper and zinc (Burkhead et al., 2009; Li et al., 2013; Hantzis et al., 2018). Hantzis et al. (2018) demonstrated that plants do implement a mechanism for iron economy that allows preferential allocation of iron to the most important functions when the demand in shoots exceeds the supply by the roots. One of the iron-depletion specific transcriptional changes is the down-regulation of ferritin. Here we showed that *FER1* expression in shoots accurately reflected the low iron levels in *cos* mutants (Fig. 3B). Thus, it appears that *FER1* expression is not impaired and shoot iron status is sensed properly in *cos* mutants.

In addition to mRNA sequencing of Arabidopsis shoots, we carried out sRNA sequencing of whole shoots under Fe-sufficient and deficient conditions. We noticed the presence of sRNAs that appear to emanate from the *COS* locus, and that are much more abundant in plants grown under iron deficiency. While searching for putative targets of these sRNAs, we found small regions of sequence complementarity to several important Fe-deficiency-induced genes (*bHLH38, bHLH39, bHLH100, bHLH101, bHLH105/ILR3, bHLH115, BTSL1, FSD1, IMA1, PYE* and *VIT1*). (Fig. 6). This surprising finding led us to speculate that the sRNAs might affect the expression of these putative targets.

Alternatively, *COS* might affect expression of these putative targets through its secondary structure. lncRNAs can form secondary structures that bind genes in trans and affect their expression (Chen et al., 2021; Wierzbicki et al., 2021). Within these secondary structures, lncRNAs can have domains where base-pairing probabilities are increased, raising the possibility that the ∼600 bp region of *COS* that contains complementary sequences for several iron homeostasis genes could be a biologically active region within the secondary structure of *COS* transcript. Interaction between the *COS* lncRNA and its targets could then affect the expression of these genes. Further investigation of the mechanism of *COS* lncRNA function should shed new light on the processes by which plants adapt to iron deficiency.

We particularly examined putative *COS* targets whose expression is associated with Fe-deficiency responses. Expression of some of these putative targets showed downward trends in expression in *cos* mutants, or failed to be fully induced during iron deficiency. We noted that putative target genes *ILR3/bHLH105, PYE*, and *bHLH101* display higher expression during iron sufficient growth of *cos* mutants. This is consistent with the possibility that *COS* negatively regulates their expression. However, *PYE* and *bHLH101*, which are both normally up-regulated during iron deficiency, both seem to have dampened iron deficiency responses in *cos* mutants. This may indicate a secondary effect in which gene expression is indirectly changed in *cos* mutants, perhaps owing to alterations in expression of upstream regulators like *PYE* and/or *ILR3/bHLH115*.

## 6 Author Contributions

E.L.W. conceived the original idea. A.B. carried out the experiments, analysed the data. A.B. and E.L.W. wrote the manuscript. All authors read and approved the final manuscript.

## 8 Acknowledgments

We thank Chris Phillips and Dan Jones at the UMass Morrill Greenhouses, for assistance with plant husbandry.

## 9 Supplementary Material

Supplementary Material should be uploaded separately on submission, if there are Supplementary Figures, please include the caption in the same file as the figure. Supplementary Material templates can be found in the Frontiers Word Templates file.

## 10 Data Availability Statement

The datasets generated for this study can be found in the National Center for Biotechnology Information (NCBI) GEO database accession number GSE209595 (https://www.ncbi.nlm.nih.gov/projects/geo/query/acc.cgi?acc=GSE209595).

