## Supplemental Material for "*CAN OF SPINACH*, a novel long non-coding RNA, affects iron deficiency responses in *Arabidopsis thaliana*"

Supplementary Material

### 1 Supplementary Figures and Tables

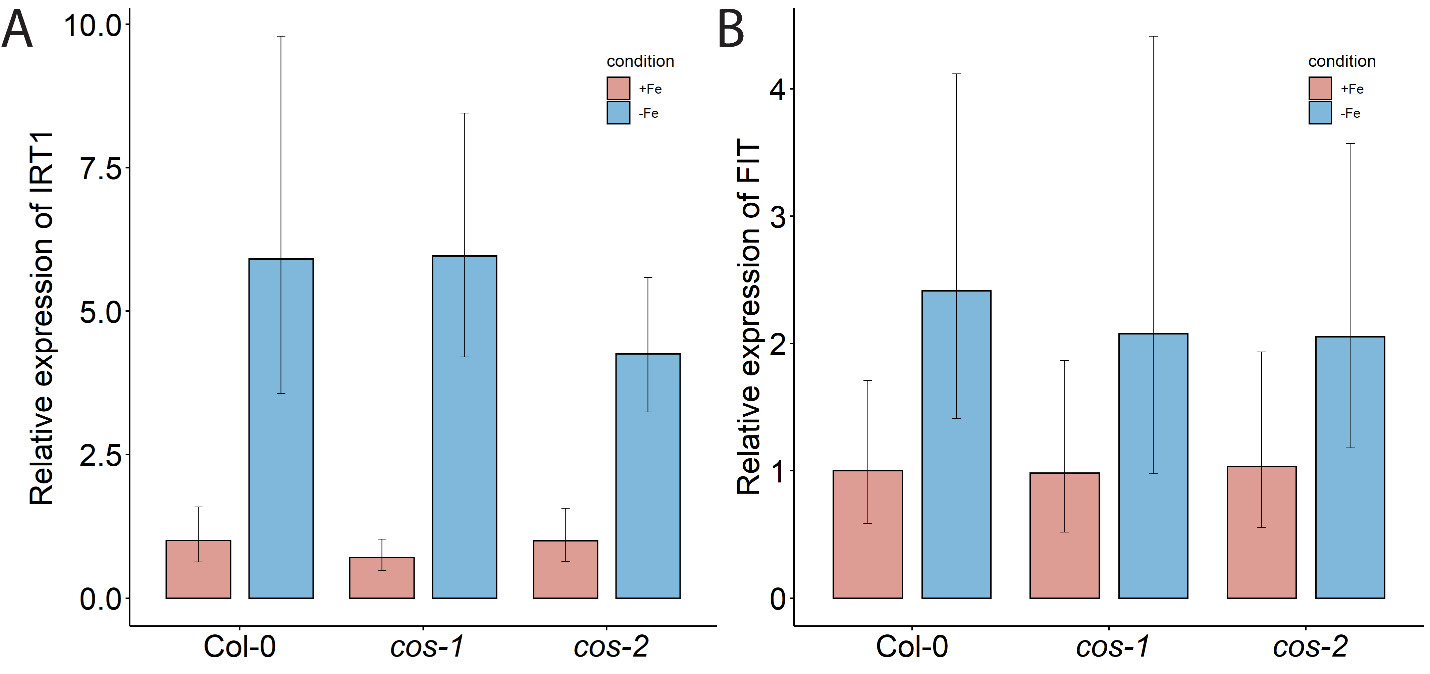

**Supplementary Figure 1.** Iron deficiency regulated gene expression in *cos* mutant roots. Relative expression levels of *IRT1* (A) and *FIT* (B) in roots of WT and *cos* mutants. Seedlings were grown on 1/2X MS agar with 50 μM Fe-EDTA for 14 d followed by a growth period of 3 d either on Fe-sufficient or Fe-deficient MS agar medium. Bars represent means ± SD of relative transcript levels normalized to *ACT2* (n=3).

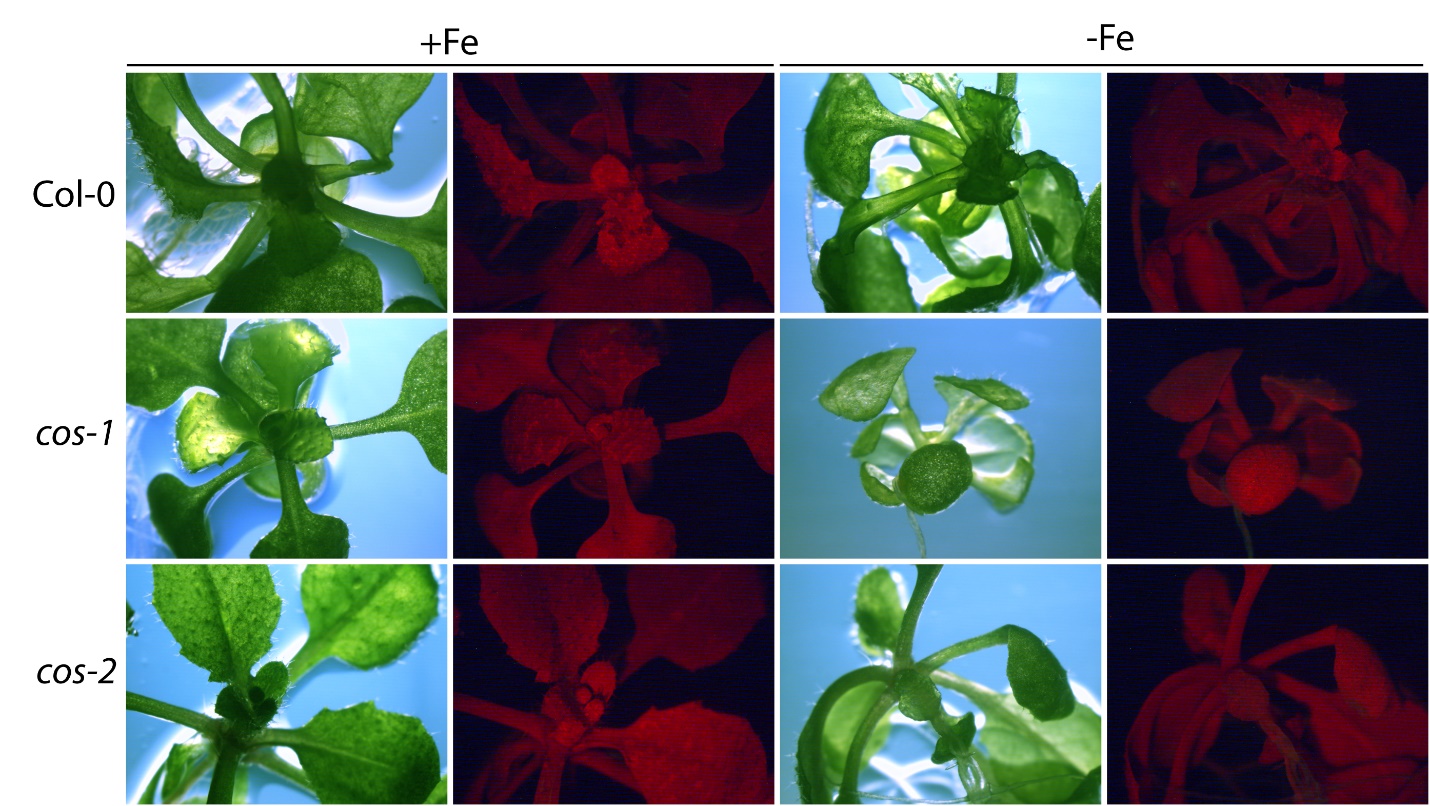

**Supplementary Figure 2.** Negative controls for SOSG treatment. Seedlings were grown on 1/2X MS agar with 50 μM Fe-EDTA for 14 d followed by a growth period of 3 d either on Fe-sufficient or Fe-deficient MS agar medium. WT and *cos* mutants were infiltrated with 50 mM phosphate buffer (pH 7.0), incubated under high light for 30 minutes and analyzed simultaneously with negative controls.

**Supplementary Table 1.** Sequences of primers used in this study

| Name | Sequence (5’ → 3’) | Reference |
| --- | --- | --- |
| COS-R1 | GGCTTCTTCTCATCCAGTTTACAAGC | This study |
| COS-F1 | GGCAACGAAACTTTCTGGACC | This study |
| COS-F2 | ACTCTCTACTTAACCATCATGCCCC | This study |
| COS-R3 | CGGTTGGATTGTTGGTTTCTGCT | This study |
| qRT-COS-F (5' end) | TGGCTTCTTCTCATCCAGTTT | This study |
| qRT-COS-R (5' end) | ACGTCATCTTTTCGTCTTTGCA | This study |
| qRT-COS-F (3’ end) | AGGTGTCCTAAGAATCTATGGCT | This study |
| qRT-COS-R (3’ end) | TCATTAACCACGTAACCGACACT | This study |
| ACT2-forward | GCTGAGAGATTCAGATGCCCA | (Kumar et al., 2017) |
| ACT2-reverse | GTGGATTCCAGCAGCTTCCAT | (Kumar et al., 2017) |
| IRT1-forward | ACTTCAAACTGCGCCGGAAGAATG | (Kumar et al., 2017) |
| IRT1-reverse | AGCTTTGTTGACGCACGGTTC | (Kumar et al., 2017) |
| FER1-forward | CAACGTTGCTATGAAGGGACTAGC | (Kumar et al., 2017) |
| FER1-reverse | ACTCTTCCTCCTCTTTGGTTCTGG | (Kumar et al., 2017) |
| FIT-forward | TTTTCGCGGTATCAATCCTC | (Oh et al., 2016) |
| FIT-reverse | GGTATGTGTCCGGAGAAGGA | (Oh et al., 2016) |
| OXI1-forward | ACGACGCTAAATTGCTTGCT | (Akmakjian et al., 2021) |
| OXI1-reverse | CCGTGAAGAGACGGAAAGAG | (Akmakjian et al., 2021) |
| AT3G03810-forward | GCTTTGACAAGAGCCACCAA | (Akmakjian et al., 2021) |
| AT3G03810-reverse | CTACTGCACGTTTGCCAGACTT | (Akmakjian et al., 2021) |
| PYE-forward | CAGGACTTCCCATTTTCCAA | (Tissot et al., 2019) |
| PYE-reverse | CTTGTGTCTGGGGATCAGGT | (Tissot et al., 2019) |
| ILR3-forward | GCAACCTATTGGTGTTTCTTCTAACTC | (Tissot et al., 2019) |
| ILR3-reverse | CCAGGTTCTTTGCTAGCTTCTGA | (Tissot et al., 2019) |
| bHLH38-forward | AGCAGCAACCAAAGGCG | (Wang et al., 2007) |
| bHLH38-reverse | CCACTTGAAGATGCAAAGTGTAG | (Wang et al., 2007) |
| bHLH101-forward | CAGCTGAGAAACAAAGCAATG | (Akmakjian et al., 2021) |
| bHLH101-reverse | CAGTCTCACTTTGCAATCTCC | (Akmakjian et al., 2021) |
| IMA1-forward | ATGTCTTTTGTCGCAAACTT | (Mankotia et al., 2022) |
| IMA1-reverse | CACCACCATTCTCACTATATG | (Mankotia et al., 2022) |
| BTSL1-forward | GGCAATGAAGATGGATTTGG | (Mankotia et al., 2022) |
| BTSL1-reverse | TCATATGGAACCGTTGCTGA | (Mankotia et al., 2022) |
